# Transposons Triggered Dynamic Evolution of MKK3 Gene, a Key Regulator for Seed Dormancy in Barley

**DOI:** 10.64898/2026.03.23.713676

**Authors:** Lydia G Tressel, Ann M Caspersen, Jason G Walling, Dongying Gao

## Abstract

Barley (*Hordeum vulgare* L.) is an important crop in the world and its seed dormancy is primarily controlled by a Mitogen-Activated Protein Kinase Kinase 3 (MKK3) gene. Although kinase activity of MKK3 and its roles in barley post-domestication have been widely studied, the pre-domestication evolution of MKK3 and the spread of nondormant alleles among global barley varieties remain largely unexplored. In this study, we analyzed MKK3 sequences in barley and its wild progenitor (*H. spontaneum*) and identified two polymorphic miniature inverted-repeat transposable elements (MITEs). Comparative analyses indicated that the insertions/excision of the MITEs predated the current estimates of barley domestication. Examination of the barley pangenomes coupled with droplet digital (dd) PCR revealed extensive copy number variation of MKK3 and suggested that transposons likely drove tandem amplification of the MKK3 gene on chromosome 5H. Additionally, approximately 1-Kb MKK3 sequences were found on chromosomes 1H and 6H. Further analysis indicated that these short MKK3 sequences were captured by a CACTA transposon that also contained fragments from four other expressed genes. The acquisition of MKK3 was estimated to be between 1.9-2.5 million years ago. Together, these findings illuminate the dynamic pre-domestication evolution of the MKK3 gene and suggest three independent origins of highly nondormant barley worldwide including a unique lineage predominant in Ethiopian germplasm. This study reveals the pivotal roles of transposons in MKK3 evolution and provide helpful information for understanding the complex history of MKK3 gene in barley and also for improving preharvest sprouting (PSH) tolerant varieties under distinct natural conditions.

## INTRODUCTION

Crop domestication marked a pivotal milestone in human history, as ancient humans gradually transformed wild species into new types of plants with favorable traits such as larger non-shattering seeds, increased apical dominance, and improved quality through prolonged artificial selection (Doebley et al. 2006). Dormancy was another key trait selected against during domestication, as it delays planting and causes uneven emergence, leading to non-uniform stands and reduced crop yield (Gubler et al. 2005; Bewley et al. 2013; Rodriguez et al. 2015). Domesticated crops exhibit significantly reduced dormancy, making them well-suited for modern agriculture; however, low dormancy increases the risk of preharvest sprouting (PHS), where mature seeds germinate on the plant before harvest under warm, moist conditions.

The substantial economic impact of crop losses due to PHS has driven intensive efforts to identify the genetic underpinnings and sources of resistance to PHS (Rodriguez et al. 2015; Matilla 2024). In the United States, approximately 68% of barley production is used for malting industries (beer), and strict malt quality standards reject barley grains showing signs of PHS, resulting in significant crop losses or severely reduced returns, a challenge further exacerbated by environmental change and expansion into nontraditional growing regions (Fox and Bettenhausen 2023; Herring et al. 2018). To address this, practices including selective breeding for PHS resistance are essential to the sustainability of the crop worldwide. Several QTL associated with PHS and dormancy have been identified in the barley genome, with one high impact locus, seed dormancy 2 (SD2) located near the distal telomeric region of chromosome 5H, drawing substantial research effort (Han et al. 1996; Lin et al. 2009; Ullrich et al. 2009; Sato et al. 2016; Nakamura et al. 2016; Nagel et al. 2019; Sweeney et al. 2021).

Mitogen Activated Kinase Kinase 3 (MKK3) was first identified in a Japanese barley population as the causative gene underlying SD2 with a mutation at the N260T codon increasing dormancy with a decreased kinase activity (Nakamura et al. 2016). Subsequent studies using varieties from United States, Canada and Northern Europe identified an alternate mutation at the E165Q codon with lines harboring this mutation demonstrating increased kinase activity and stark PHS susceptibility (Vetch et al. 2020; Jorgenson et al. 2026). Malt quality traits correlate well with germination energy and the allelic effects of MKK3 can alter malt barley profiles (Woonten et al. 2005; Rooney et al. 2023). For example, malting lines with the highly nondormant E165Q mutation result in higher levels of hydrolytic enzymes (namely alpha and beta amylase) and free amino nitrogen (FAN), both characteristic of an adjunct style malt popular in the northwest growing region of United States and western Canada (Beattie et al. 2010; Rooney et al. 2023). The MKK3 locus exhibits extensive copy number variation (CNV), including cases of gene amplification up to 15 copies. Pangenome analysis revealed that these multicopy arrays often contain a mix of wild-type and mutated alleles, and lines most susceptible to PHS typically carry multiple MKK3 copies, including one or more with the highly nondormant E165Q variant (Jorgenson et al. 2026).

Recent insights into the role of the MKK3 locus in barley domestication revealed a narrative of human migration, where specific allelic and CNV combinations of MKK3 adapted to environments that supported reliable production and enhanced end-use quality (Jorgenson et al. 2026). For example, the E165Q variant, now widespread in North American germplasm and largely responsible for the success of adjunct-style malts but also contributes to rising PHS problems (Rooney et al. 2023). This allele likely originated in Northern Europe and Scandinavia, where cool growing seasons favor dormancy and minimize PHS risk, thereby varieties with rapid, uniform germination would have been advantageous (Sweeney et al. 2021; Jorgenson et al. 2026). ‘Bere,’ one of the oldest known barley varieties and leveraged in Scotland for thousands of years, harbors the E165Q mutation and is surmised as a potential early source of this nondormant genotype as seed stocks spread through migration and expanding trade networks (Kjær et al. 2025). In contrast, varieties with single copy MKK3 and dormant alleles appear to have become established in regions characterized by high precipitation and humidity near harvest, where stronger selection against PHS would have favored dormancy (Nakamura et al. 2016; Jorgenson et al. 2026). While the post-domestication history of MKK3 and its impact on barley production provides a clearer understanding of the genetics and population diversity of MKK3 underlying worldwide barley production, the evolutionary dynamics of MKK3 before domestication remain poorly understood.

Transposons are ubiquitous elements of plant genomes and are classified into two Classes: Class I RNA retrotransposons and Class II DNA transposons. Retrotransposons are stable insertions since they remain positioned in the host genome at the place of insertion and do not excise once they are established. In contrast, the insertions of DNA transposons can be unstable as many of them may excise and frequently leave behind shorter DNA sequences at the integration sites called footprints (Scott et al. 1996). Except for a few superfamilies such as Helitrons, nearly all transposons generate target site duplications (TSDs) when they insert into new sites, but the TSDs may vary in size between different superfamilies (Wicker et al. 2003). Miniature inverted-repeat transposable elements (MITEs) are a specialized type of DNA transposon that contains terminal inverted repeats (TIRs) and are flanked by TSDs, the typical features of many DNA transposon superfamilies. However, they are very small (usually 100-500 bp) and do not encode transposase (Bureau and Wessler 1992; Han and Wessler 2010). Transposons have contributed to chromosome stability, centromere/telomere formation (Wong and Choo 2004), and their insertions can alter gene expression and induce genetic variability essential to natural selection and host fitness (Barrón et al. 2014). By comparing their movement and accumulation between related organisms, transposons may also serve as “molecular fossils” for tracking evolutionary history of previous genomic events (Le Rouzic et al. 2013).

In this study, we analyze the MKK3 gene sequences and our primary objectives include: 1) reveal MKK3 sequence variations in barley and its wild relatives; 2) understand evolutionary history of MKK3, especially for pre-domestication evolution; and 3) test the hypothesis of single origin of low dormant barley in North America. Our analyses provide new insights into MKK3 evolution and the origins of nondormant barley cultivars across the world.

## RESULTS

### MITEs-mediated MKK3 gene variation

Using BLASTN with the MKK3 sequence of barley variety Steptoe (**Table S1**), hereafter called reference MKK3 gene, we observed two insertions or deletions (InDels) among the published MKK3 genes (Nakamura et al. 2016). One InDel was 769-bp and located in the second intron, and another was 338-bp and located in the fifth intron (**Figure 1A**). We manually inspected these two InDels and found that the larger InDel contains a 160-bp MITE designated Hvu_MITE1 and the smaller InDel contains a 335-bp MITE designated Hvu_MITE2. In addition, another copy of Hvu_MITE1 was also detected in the seventh intron but it is shared by all MKK3 genes. To distinguish the two copies of Hvu_MITE1, the MITE located in the second and seventh introns were designated Hvu_MITE1a and Hvu_MITE1b (**Figure 1B**). Based on the presence/absence of two MITE families, the sequenced MKK3 genes were further divided into three groups (T1, T2 and T3) for downstream analysis and defined as: T1 that only contains Hvu_MITE1b, T2 that contains both Hvu_MITE1a and Hvu_MITE1b, and T3 that contains all three MITEs (**Table S1**).

**Figure 1.**
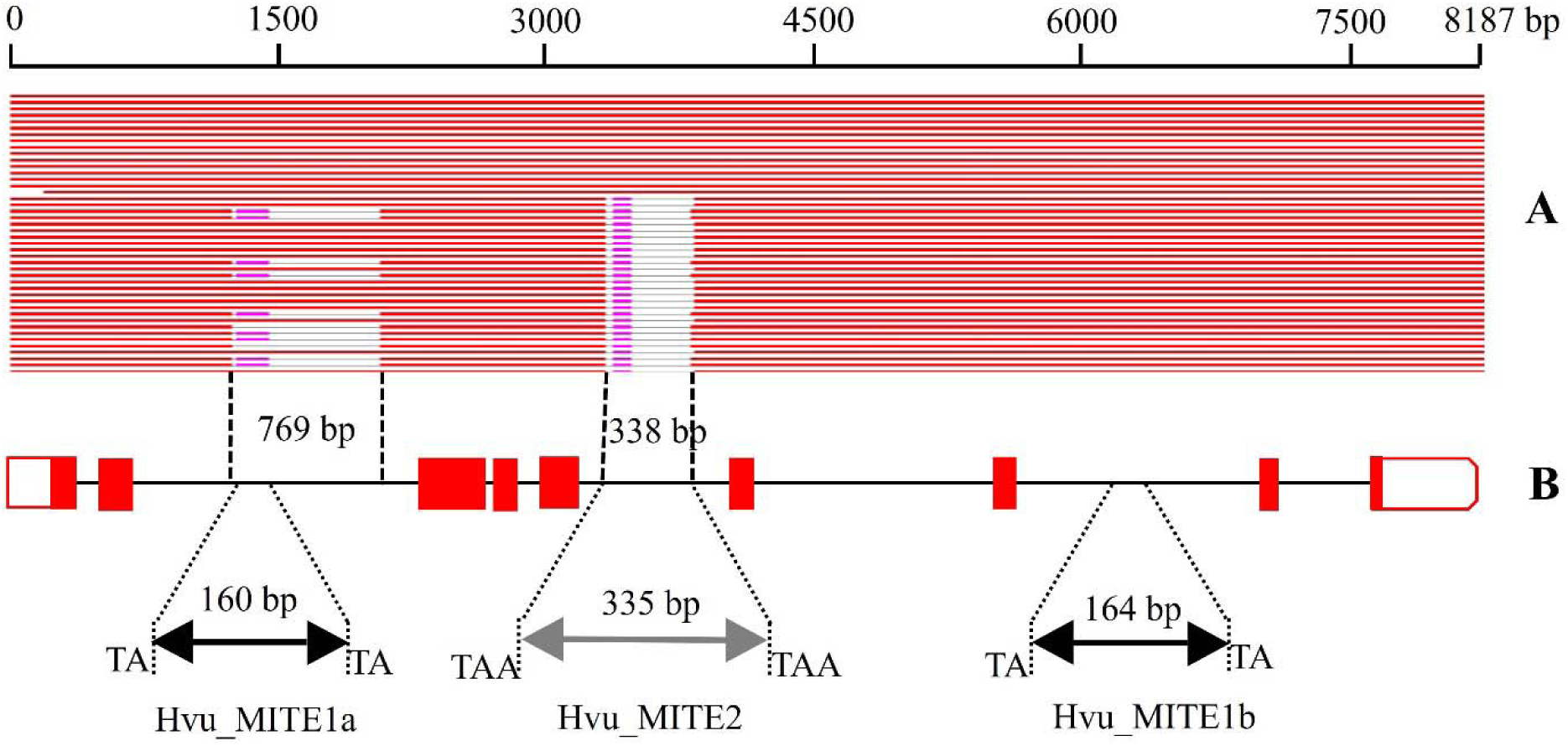
A. BLASTN graphics with the MKK3 gene in Steptoe. B. The targeted regions of three MITEs in the MKK3 gene. The red boxes and black line denote exons and introns of MKK3. The black and grey triangles represent the TIRs of MITEs. The purple hits represent unfaithful sequence alignment with Hvu_MITE1b in 769-bp and a short tandem in 338-bp InDel.

To gain insight into the insertion times of the three MITEs, the MKK3 genes from three barley varieties and three wild barley accessions were aligned. Additionally, the alignment also included MKK3 homologues from common wheat (*Triticum aestivum* L.) and bulbous barley (*H. bulbosum*) which represents the closest wild relative to barley and its wild progenitor. No Hvu_MITE1a was detected in both wheat or bulbous barley, however, both homologs shared flanking sequences with two barley varieties (Steptoe and Harrington) and two wild barley accessions (ICWB180001 and ICWB181450) (**Figure S1A**), thus Hvu_MITE1a inserted into the barley genome after the split from bulbous barley which is estimated to have occurred about 4.5 million years ago (MYA; Brassac and Blattner 2025), after which the transposon appears to excised itself, carrying it with some flanking sequence or the MITE and the flanking sequence has been eliminated by genomic selection pressure. The homologous sequences in both wheat and bulbous barley contain neither Hvu_MITE2 nor Hvu_MITE1b, and the MKK3 genes in barley contain the MITEs plus the TSD (TAA or TA; **Figure S1B, C**), thus, these two MITEs also inserted into barley after the split from bulbous barley but remained static in their position after the insertion event. Since three MITEs were found in the MKK3 genes from both barley and wild progenitor, they should integrate into MKK3 prior to barley domestication.

### Duplication of MKK3 sequences

Our previously Sanger sequencing of the MKK3 coding region in PHS susceptible malting barley varieties often resulted in a detected double peak at the 2,506^th^ nucleotide of reference MKK3 gene or the MKK3 Q165 which is related to exchange of a Glutamic acid (E) to Glutamine (Q) at position 165 (Vetch et al. 2020) (**Figure S2A-C**) that could be interpreted as heterozygosity at the MKK3 locus (Sweeney et al. 2021) but would counter the presumed homozygosity of inbred barley varieties. To detect if multiple MKK3 sequences are present in other barley genomes, the reference MKK3 gene was used to search against 86 barley genomes published by eight research teams (Liu et al. 2020; Zeng et al. 2020; Xu et al. 2021; Jiang et al. 2022; Hu et al. 2023; Pan et al. 2023; Clare et al. 2024; Jayakodi et al. 2024) (**Table S2**). Among the 62 cultivated barley genomes, 55 contain only a single copy of the complete MKK3 on chromosome 5H, and seven contain multiple copies of complete MKK3 including two in Morex, OUN333, HOR2180 and Hockett, three in HOR7552, four in HOR12184 and five in 10TJ18, and all these duplicated copies were tandemly organized on chromosome 5H. However, no duplicated complete MKK3s were found in the 24 wild barley, so all of them contained a single copy of MKK3 on chromosome 5H (**Table S2**).

To confirm the copy number variation of MKK3, droplet digital (dd) PCR was performed for 40 cultivated and wild barley genomes including ten for which the genomes have been published. One to four copies of the MKK3 gene were detected in the 40 genotypes indicating gene duplications in some barley varieties (**Table S3**). The ddPCR results were consistent with genomic analysis of the MKK3 gene as well as the published results of MKK3 (Jorgenson et al. 2026) in seven sequenced genomes including Morex, Golden promise and five wild barley accessions, but were different for three genomes. ddPCR detected three copies of MKK3 in both AAC Synergy and Hockett genomes. However, only one and two copies of the complete MKK3 gene were found in the genomes of AAC Synergy and Hockett suggesting the possibility that some MKK3 sequences may have been misassembled. Importantly, the MKK3 genes in these 86 sequenced genomes can also be grouped into three groups based on the polymorphic MITEs (**Table S2**). To validate the presence/absence of the MITEs, PCR analyses were conducted using two pairs of PCR primers that targeted the flanking region of the 769-bp InDel and Hvu_MITE2, respectively. Expected sizes of amplicons were observed (**Figure S3**) and further confirmed the polymorphisms of the two MITEs.

### Phylogeny of MKK3 genes

To understand the evolutionary relationships of MKK3 genes, 179 complete MKK3 sequences from barley and its wild progenitor were used for phylogenetic analysis with maximum likelihood methods. The homologue of barley MKK3 found in *H. bulbosum* was also included and used as the outgroup. The resulting phylogeny grouped the 179 MKK3 genes into three major clades. All T1-group MKK3 genes were grouped into Clade II, all T2-group MKK3 genes into Clade III, and all MKK3 genes fell into the two subclades of Clade IV (**Figure 2**). To rule out the possibility that the insertions/excision of MITEs in MKK3s may affect alignment accuracy and generate phylogenetic noise, the alignment regions containing 769-bp InDel and 338-bp InDel were removed, and the shared sequences of 179 MKK3 genes were used to build another phylogenetic tree with the same pipeline. A similar phylogeny was obtained, T1, T2 and T3 groups of MKK3 genes fell into Clade II, III, and IV, respectively (**Figure S4**). Regardless of the input sequences used, the MKK3 genes in cultivated and wild barley with the same transposon patterns were grouped into the same clades. The MKK3 genes in the wild progenitor were not grouped into a single lineage and separated from the MKK3s in cultivated barley further suggested that the events of insertions and excision of the MITEs occurred prior to barley domestication. Both phylogenetic trees indicated that the branches of T1 group MKK3 were closest to the outgroup than the other two.

**Figure 2.**
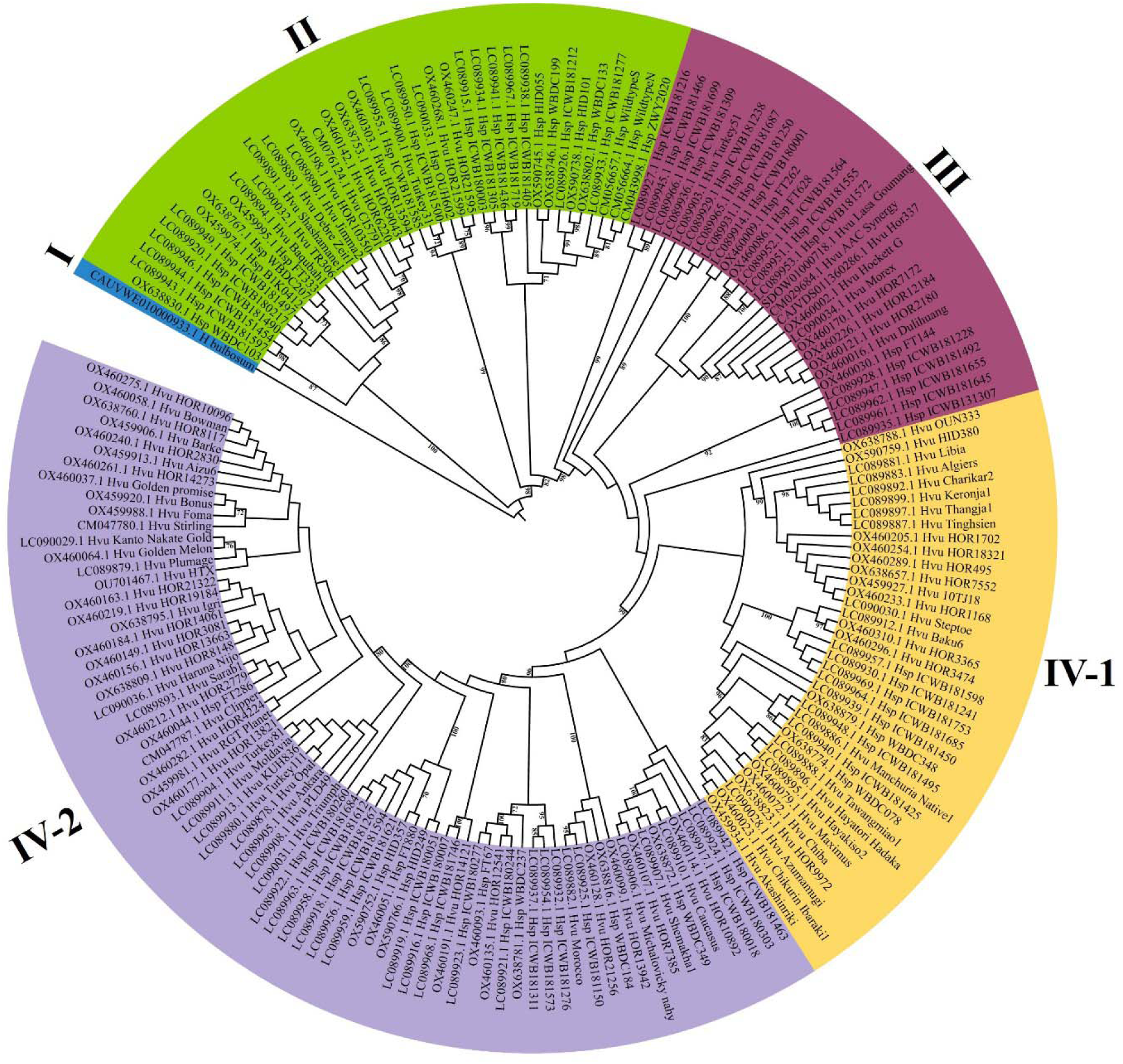
Phylogenetic tree of 179 complete MKK3 gene sequences in cultivated and wild barley built with maximum likelihood methods. The bootstrap values of >70% are labeled. The three letters between the GenBank accession number and the cultivar/wild accession represent cultivated (Hvu) and wild progenitor (Hsp) barley respectively.

### Germination rates and expression of different MKK3 groups

We leveraged published data of barley seed germination (Nakamura et al. 2016) to investigate whether the MITEs in MKK3 confer a demonstrable impact on seed germination. Among the 274 barley cultivars and 150 wild barley accessions, 236 and 118 have information on sequence types of MKK3 that were used to define the MKK3 gene group for each genotype based on their sequences (**Table S4**). 94 wild accessions showed lower germination rates ranging from 0 to 22% and suggested that they generated seeds with strong dormancy. However, 10 wild accessions exhibited moderate or higher germination rates ranging from 50% in ICWB181573 to 79% in ICWB181753. These accessions included two with T1, three with T2 and five with T3 type MKK3. Among 186 cultivated genotypes with T3-type MKK3, the germination rates of 40 cultivars were lower than 25%, and 111 showed higher germination rates ranging from 60% to 100%. For the 12 barley varieties with T2-type MKK3, nine showed high germination rates of over 89% whereas Turkey 51 and Bursa exhibited extremely low germination rates of less than 4%. For the 36 cultivars with T1-type MKK3, 33 showed high germination rates of over 86% but Turkey 31 had a germination rate of 0.41%. Overall, dramatic variations of seed germination rates were observed within the domesticated barley varieties as the cultivars with extremely low and high germination rates were found for each MKK3 group (**Figure 3A**).

**Figure 3.**
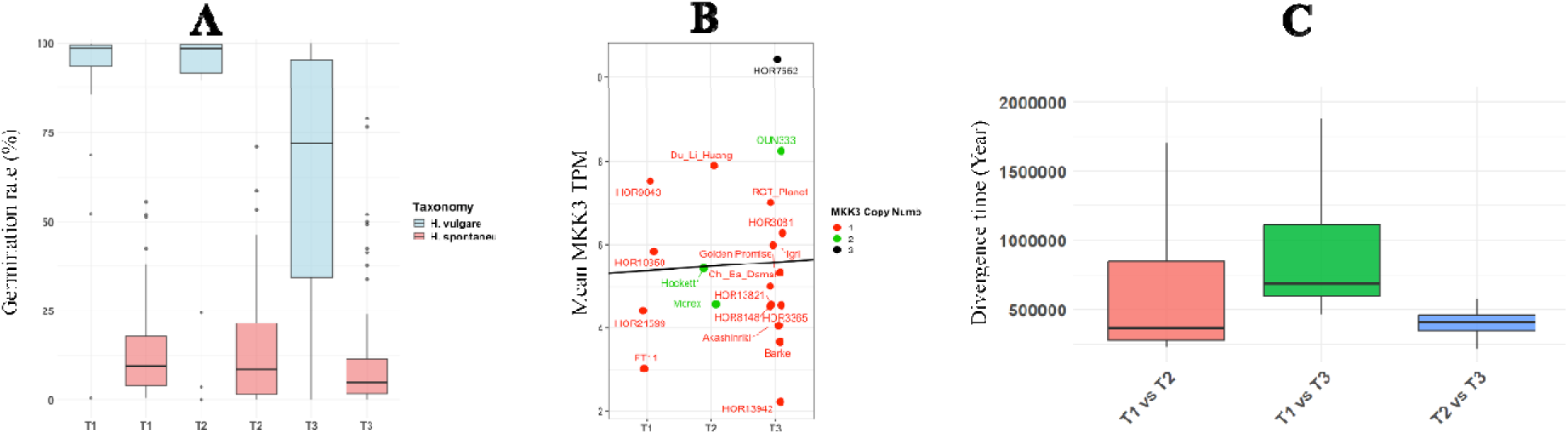
A. Boxplots of germination rates of 236 cultivated and 118 wild barley genotypes. The germination rates were calculated based on the published results (Nakamura et al. 2016). **B. MKK3 gene expression in the embryonic tissue of 19 cultivated and one wild barley. C. Divergence estimation between three established MKK3 groups.**

We further used the barley pan-transcriptome data (Guo et al. 2025) to query whether the presence of MITEs can affect expression levels of MKK3. We limited our analysis of the RNA-seq data to that generated from the embryonic tissue since it has inherently higher MKK3 transcript levels than other tissues (Jørgensen et al. 2026). Variable abundance of transcripts was found between the 20 genotypes, however, the total transcripts per million (TPMs) were less than 9 TPMs in 19 genotypes, suggesting low to modest levels of gene expression (**Figure 3B**). The highest TPM was observed in HOR7552, and contains three copies of the MKK3 gene. No significant differences in MKK3 gene expression were found between T1, T2 and T3 groups (*P*-value > 0.05), although not surprising as all three MITEs occupy intronic regions of MKK3.

To gain insight into the divergence of three MKK3 groups, all MKK3 genes in one group were used to conduct all-against-all comparisons with the MKK3s in other two groups to calculate the synonymous rates. The mean divergence time of MKK3s was 940,933 years between T1 and T3 group, 627,758 years between T1 and T2 group, and 400,145 years between T2 and T3 groups, respectively (**Figure 3C**). Thus, based on the resulting grouping, the T1 group may be the older MKK3 whereas the T3 group likely evolved more recently.

### Repeat-related MKK3 gene duplication

The CNV of the MKK3 gene was previously reported (Jørgensen et al. 2026), and our ddPCR analysis confirmed these CNV values while also provided estimates for additional varieties (**Table S3**). However, the genomic basis and timing of the MKK3 duplication were not previously examined. To address this, we extracted sequences of the duplicated MKK3 genes along with 11-kb of flanking regions (5.5-kb on each side) from chromosome 5H and compared them across seven genomes containing multiple MKK3 copies. We also analyzed MKK3 and its flanking regions in two wild species (WBDC199 and WBDC349) that carry a single-copy gene. A 3.8-Kb mutator transposon was found in the 1-Kb region of upstream MKK3 gene in both WBDC199 and WBDC349. Furthermore, a truncated Helitron transposon and ∼840-bp tandem repeat sequence were detected in ∼2-Kb region downstream the MKK3 gene. Impressively, these repeats were also detected in all duplicated genes in seven barley genomes including HOR12184 in which the duplicated MKK3s were in reverse orientation, and they showed identical organization patterns with MKK3 (**Figure 4**). This suggests MKK3 was duplicated together with these repeats. Although the exact mechanism remains unclear, the presence of repeats strongly implies their involvement in the duplication process. Furthermore, identical MITEs polymorphisms across copies indicate duplication occurred after divergence of the three MKK3 groups.

**Figure 4.**
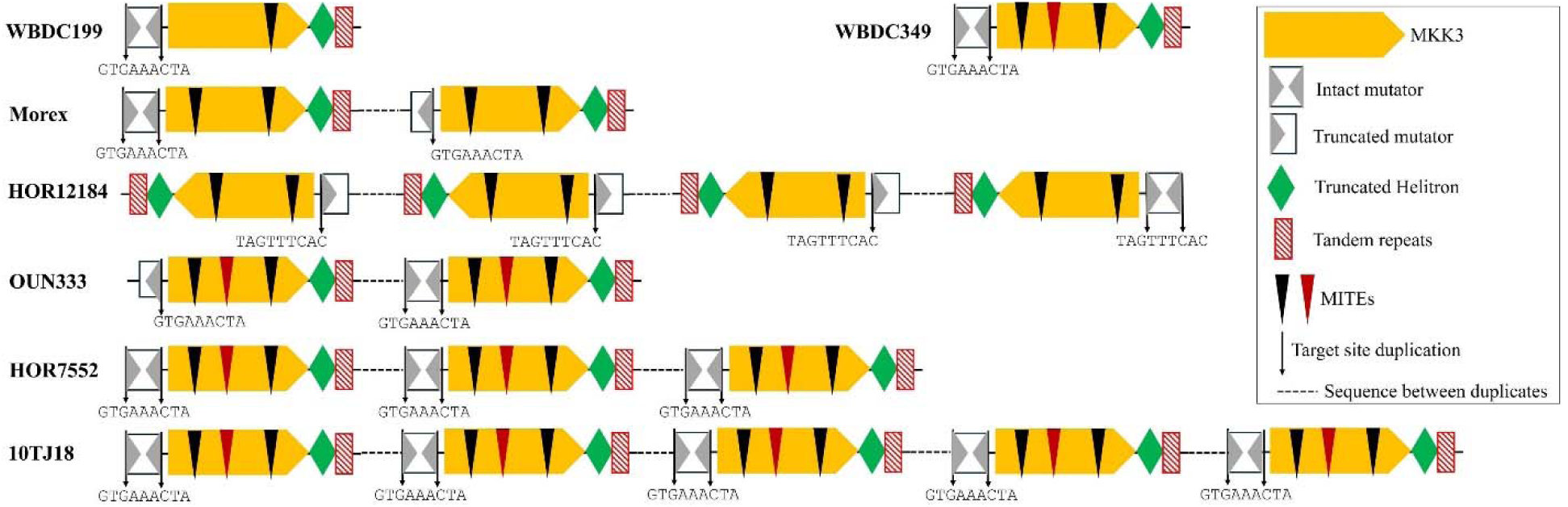
MKK3 gene and flanking repeats in two wild barley accessions (WBDC199 and WBDC349) with a single complete MKK3 on 5H and five barley cultivars with two or more complete MKK3 on 5H. The 9-bp TSD of the Mutator transposon is shown, and the MKK3s in HOR12184 were in reverse orientation.

The two copies in Morex share 100% sequence identity suggesting this duplication was of recent origin. The two MKK3s in HOR2180 also share 100% sequence identity to most of the gene, except for a variable region between 6790-7140 bp that may be caused by inversion or errors in sequence assembly. The duplicated MKK3s in Hockett, HOR7552, HOR12184 and 10TJ18 either showed 100% sequence identity or only contained a single point mutation at different positions in the four genomes including G to C at 2506^th^ in Hockett, C to T in 4572^th^ in HOR7552, G to A in 4772^th^ in HOR12184 and T to C at 5915^th^ in 10TJ18. However, multiple point mutations and InDels ranging from 1-bp to 17-bp were found in duplicated MKK3s in OUN333. The duplicated MKK3s were aligned and used to calculate synonymous rates. The divergence time of duplicated MKK3s in Morex and Hockett was 0 years and 9,384 years, respectively. Similar estimation was also obtained in HOR7552, HOR12184 and 10TJ18 that contain more than three MKK3s, and the divergence time ranged from 0 to 9,384 years depending on duplicated pairs for calculations. These results indicate that gene duplication events in the five genotypes were relatively recent, likely occurring after barley domestication in the Fertile Crescent about 10,000 years ago (Badr et al. 2000). In contrast, the divergence time between the two MKK3 copies in OUN333 is estimated at approximately 264,615 years, indicating a more ancestral origin than the time frame of barley domestication. It is important to note that molecular clock-based estimates have limitations in accurately dating very recent sequence variations since a single point mutation can emerge at any time within 9,384 years.

### Capture and movement of MKK3 gene

In addition to the complete MKK3 gene on chromosome 5H, a significant hit (E-value = 0) was detected on chromosome 6H in all 86 sequenced genomes except ‘Lasa Goumang’ for which the contigs were not assembled into chromosomes. The hits on chromosome 6H ranged in size from 1,001-bp to 1,011-bp and matched the 1,019-bp region (from 76^th^ to 1094^th^) of reference MKK3. Another significant hit (E-value = 0) ranging in size from 1,022-bp to 1,044-bp was found on chromosome 1H in 12 cultivated and 9 wild barley genomes (**Table S2**). These hits also matched the 1,019-bp region (76^th^ - 1094^th^) of reference MKK3. To distinguish the two fragmentary MKK3s, the MKK3 sequence in 1H and 6H was named MKK3-f1H and MKK3-f6H, respectively.

To investigate the evolution of fragmentary MKK3s and assess whether the same repetitive sequences flanking the complete MKK3 on 5H contributed to their duplication, we analyzed MKK3-f1H and MKK3-f6H along with 11-kb of flanking sequences and compared them to the complete MKK3 region and its flanking sequence in the HOR10892 genome. The flanking regions of MKK3-f1H and MKK3-f6H showed no similarity to that of the complete MKK3 sequence but they shared ∼95% identity with each other. Manual inspection revealed that both flanking regions contain two structural hallmarks of CACTA superfamily transposons: 1) TIRs starting with “CACTA” and ending with “TAGTG”, and 2) 3-bp TSDs. These features indicate that MKK3-f1H and MKK3-f6H are harbored by two copies of the same CACTA transposon family designated CAT250.

The 5,875-bp transposon located on 1H (CAT250-1H) and the 5,698-bp element on 6H (CAT250-6H) are non-autonomous because they do not encode functional transposase. To elucidate the molecular function of the internal regions between their TIRs, CAT250-1H and CAT250-6H were queried against the barley expressed sequence tags (ESTs) in GenBank. The analysis results in over 100 significant hits (E-value < 1e-73). BLASTX using the four top hits (E-value = 0, > 500-bp) indicated these sequences were fragments of different functional genes. Subsequent searches against the annotated genes in the Morex reference genome indicated that CAT250-1H and CAT250-6H contain six fragments from five genes (**Figure 5A**):

- *HORVU.MOREX.PROJ.2HG00181110.1* encoding UDP-glycosyltransferase,
- *HORVU.MOREX.PROJ.5HG00434360.1* encoding dihydroflavonol-4-reductase,
- *HORVU.MOREX.PROJ.6HG00476760.1* encoding disulfide isomerase,
- *HORVU.MOREX.PROJ.5HG00773660.1* encoding phosphate translocator
- *HORVUMOREX.PROJ.5HG00474740.1-*MKK3.

**Figure 5.**
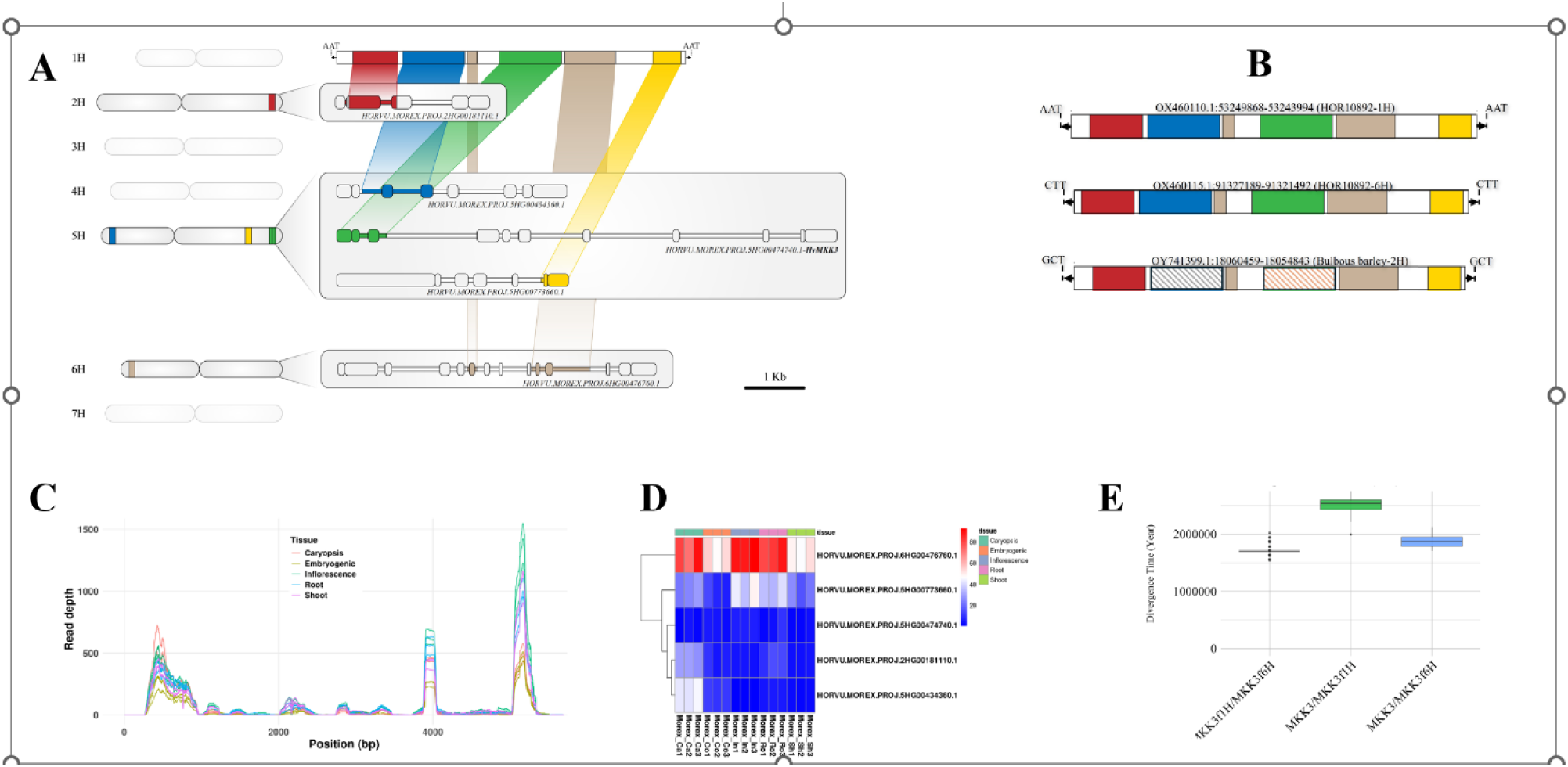
A. Fragments captured by CAT250 transposon from five genes on 2H, 5H and 6H. B. Two CAT250 copies in the barley cultivar HOR10892 and its homologous transposon in *H. bulbosum*. The red, grey and yellow boxes indicate the shared fragments acquired from three genes before the divergence between barley and *H. bulbosum*. The blue and green boxes indicate the captured gene fragments in barley after the divergence. The boxes with stripes in *H. bulbosum* indicate the unshared sequences. The white boxes indicate shared transposon sequences between HOR10892 and *H. bulbosum*. Arrows delineate TIRs of CAT250 and the 3-bp TSDs. **C. Mapped RNA-seq reads on the CAT250 transposon in the Morex genome. D. Heat map of five annotated genes in Morex.** The fragments of these five genes were captured by the CAT250 transposon. **E. Divergence time estimation between MKK3, MKK3-f1H and MKK3-f6H**.

Thus, MKK3-f1H and MKK3-f6H represent MKK3 fragments captured by CAT250 transposon. Since distinct TSDs were identified for CAT250-1H and CAT250-6H (**Figure 5B**), transposition of CAT250 likely occurred after the gene acquisitions. To further assess expression, we analyzed the RNA-seq data in Morex (Guo et al. 2025) and found that the gene fragments within CAT250 were targeted by varying numbers of sequencing reads (**Figure 5C**). Additional analysis revealed that the five annotated genes were expressed across five different tissue types in Morex (**Figure 5D**).

To understand the history of gene acquisitions in CAT250, we employed two complementary approaches: comparative genomic analysis and molecular clock estimation. First, CAT250-1H and CAT250-6H were queried against the bulbous barley genome. No related transposon was detected on 1H and 6H in bulbous barley, however, a homologous transposon was found on 2H and flanked by different TSDs. Impressively, it shared the fragments of *HORVU.MOREX.PROJ.2HG00181110.1*, *HORVU.MOREX.PROJ.6HG00476760.1* and *HORVU.MOREX.PROJ.5HG00773660.1* with CAT250-1H and CAT250-6H but lacks the fragments of *HORVU.MOREX.PROJ.5HG00434360.1* and MKK3 (**Figure 5B**). Thus, the CAT250 transposon likely captured the three gene fragments before the divergence between barley and bulbous barley but didn’t acquire fragments of MKK3 and *HORVU.MOREX.PROJ.5HG00434360.1* until after their divergence. Second, MKK3-f1H, MKK3-f6H, and the full-length MKK3 on chromosome 5H from each of the 86 sequenced barley genomes were aligned to estimate pairwise divergence rates. For the genomes lacking MKK3-f1H (**Table S2**), only MKK3-f6H and the full-length MKK3 were compared, yielding mean divergence times of approximately 2.54 MY for MKK3/MKK3-f1H and 1.87 MY for MKK3/MKK3-f6H. Thus, the estimates suggest that the MKK3 gene captured by CAT250 likely occurred between 1.9-2.5 MYA (**Figure 5E**). Taken together, the two different approaches produced consistent estimates, indicating that MKK3 acquisition occurred after the divergence between barley and bulbous barley. Furthermore, the mean divergence time between MKK3f1H/MKK3f6H was about 1.7 MY, providing a preliminary indication of the timing of CAT250 transposition, however additional data is needed to improve the precision of this estimate.

### PCR detection of polymorphic MITEs in Bere and Northern European cultivars

The Bere barley was proposed to be the ancestral donor of MKK3Q165 that is widespread in North American germplasm (Sweeney et al. 2021; Jorgenson et al. 2026). However, the genomic information of Bere accessions is limited and only Bere Unst has been sequenced (Kjær et al. 2025). Its 7797-bp MKK3 contains both Hvu_MITE1a and Hvu_MITE1b but lacks Hvu_MITE2 suggesting it belongs to T2-type MKK3. We also conducted PCR analysis in 29 barley varieties including 13 unsequenced from Northern Europe and the two Bere accessions (Bere and Scot Bere). The amplicon with an expected larger size was obtained in all tested genotypes with the primers targeting the flanking region of the larger InDel and suggesting that they contain Hvu_MITE1a. For the PCR amplification with the primers targeting the flanking region of Hvu_MITE2, expected larger amplicon was obtained in Bently, Betzes and Steptoe indicating the MKK3 in these three genotypes contain Hvu_MITE2, and the smaller size product was observed in the remaining 26 genotypes (**Table S5**) suggesting their MKK3 lack Hvu_MITE2. Thus, our PCR results indicated that two Bere accessions (Bere and Scot Bere), 13 Northern European cultivars and 10 North American varieties contain T2-type MKK3 suggesting their common origin.

## DISCUSSION

### Pre-domestication evolution of MKK3

Extensive research has characterized allelic variation within the barley MKK3 locus associated with the post-domestication reduction in seed dormancy; however, our current understanding of sequence divergence prior to barley domestication is limited (Nakamura et al. 2016; Vetch et al. 2020; Jorgenson et al. 2026). We leverage published MKK3 sequences to conduct comparative analysis between the genomes of barley and other species, and provide the first insights into the pre-domestication evolution of MKK3. We found that the ancestral MKK3 lacked transposons, and that the insertions/excision/elimination of MITEs occurred after the divergence between bulbous barley and barley wild progenitor resulting in three distinct MKK3 gene types. Our data also reveals that fragmentary MKK3 sequences were acquired, mobilized and duplicated by CACTA transposon suggesting that the gene capture events likely persisted for at least 2.6 MY (from more than 4.5 to 1.9 MYA). Striking copy number variation at the MKK3 locus was reported and our analysis of an additional 10 published genomes that were not included in the barley pangenome (Jayakodi et al. 2024), together with ddPCR of other genotypes including the seven wild accessions, further confirms the CNV landscape of MKK3 (Jorgenson et al. 2026). The presence of shared repeats flanking duplicated copies on 5H strongly points to a transposon-driven mechanism underlying MKK3 amplification, a hypothesis that certainly warrants further investigation. Comparative analysis of the duplicated MKK3s indicated that gene duplication occurred either before or after the domestication of barley. Notably, of the 86 analyzed genomes, we only identified duplicated copies of MKK3 on 5H in seven cultivated barley genomes, and while this observation is provocative, it does not conclusively indicate that duplication was confined to domesticated barley. Genes may be duplicated in wild species but are generally rendered inactive due to the actions of purifying selection (or genetic drift) that allows for accumulation of genetic mutations forming inactive pseudogenes or gene loss in wild species (Lynch and Conery 2000; Qian et al. 2010). However, the point mutations at E165Q and N260T appear to have arisen after domestication as all wild barley accessions evaluated harbor the same alleles at the two loci. Overall, MKK3 may represent one of, if not the most, dynamic genes controlling important agronomic traits in the barley as it has undergone an extraordinary series of evolutionary events over the past 4.5 MY, including insertions/excision or elimination of MITEs, transposon-mediated gene acquisition and mobilization, gene duplication and point mutations (**Figure 6**). These events generated divergent MKK3 gene sequences that underpin its critical role in regulating seed dormancy and stress signaling pathway and provide a valuable source of plasticity enabling both natural and artificial selection and reveal an opportunity to continue to push barley cultivar adaptation to suit increasingly diverse growing environments (Zhang et al. 2012; Goyal et al. 2018).

**Figure 6.**
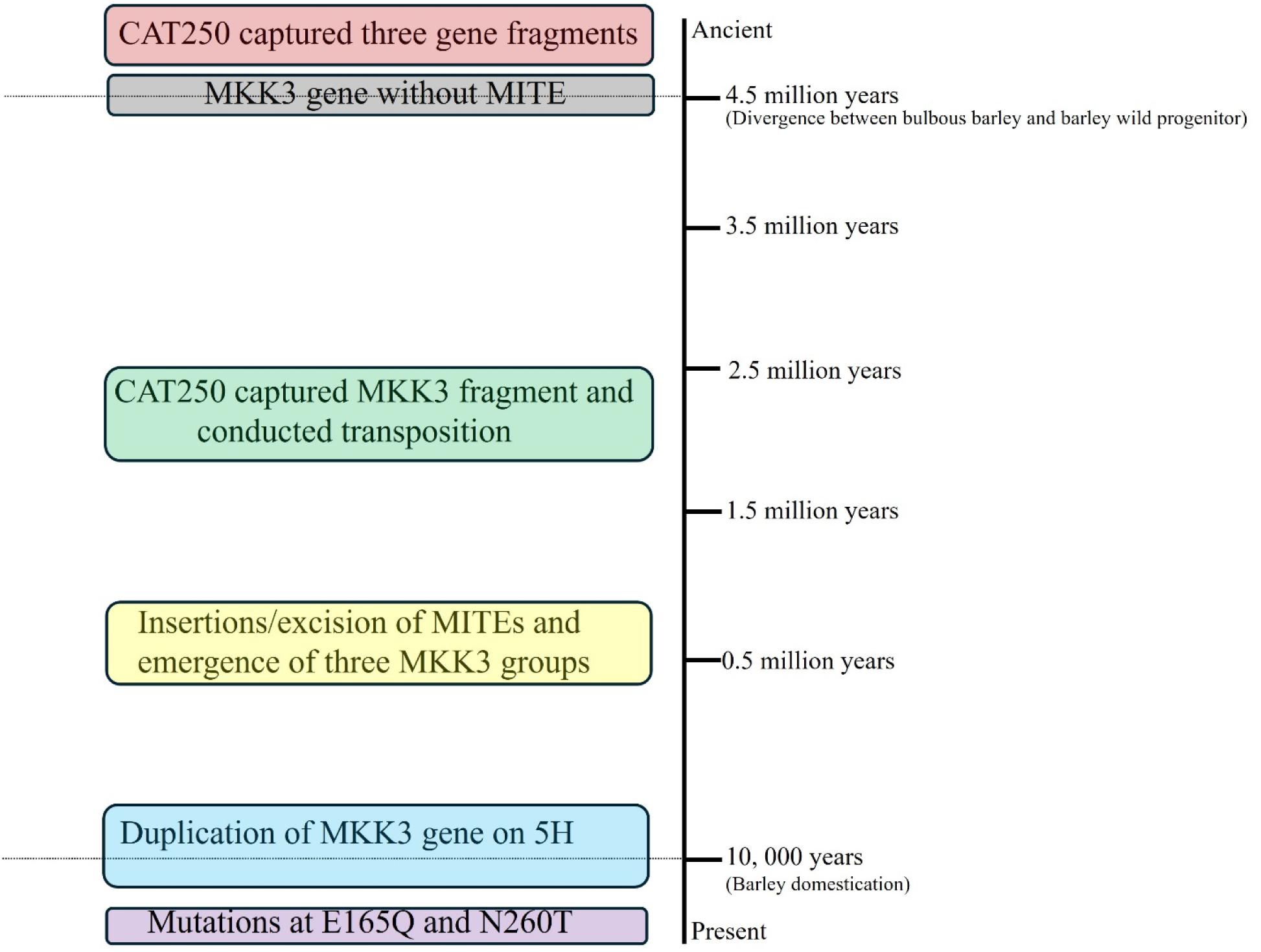
The major evolutionary events of MKK3 in the past 4.5 million years. The broken lines indicate the times of divergence between barley and *H. bulbosum* and barley domestication.

### Multiple origins of highly nondormant barley worldwide

Previous studies suggested that the highly nondormant trait, common in much of North American barley germplasm, likely originated from the ancestral donors in Northern Europe or the Bere barley accessions in Scotland where the environment favors nondormant alleles (Sweeney et al. 2021; Jorgenson et al. 2026). The ancestral allele at N260T of the PHS tolerant cultivar ‘Azumamugi’, which may be associated with nondormancy in the cultivar ‘Kanto Nakate Gold’ (KNG), is present in the wild progenitor as well as the barley cultivars collected from East Asia and other regions (Nakamura et al. 2016). However, whether the highly nondormant E165Q mutation shares the same origin with KNG and other cultivars in East Asia is still unclear. Our analysis found that MKK3 in North American malting barley varieties such as Harrington, Hockett, and AAC Synergy belongs to the T2 group that also includes the gene in Bere Unst, Bere, Scot Bere and the 13 cultivars collected in Northern Europe. However, the MKK3 gene in KNG and several other East Asian cultivars belongs to the T3 group which represents the most widespread MKK3 lineage and found in the highly nondormant cultivars across Europe, Asia, Africa, and North America (**Table S4**) as well as the Australian cultivar Clipper (Hu et al. 2023). This suggests that highly nondormant barley genotypes in North America may trace back to two donors: cultivars with T2-type MKK3 in Northern Europe or Scotland (Vetch et al. 2020; Jorgenson et al. 2026), and cultivars with T3-type MKK3. The nondormant Japanese cultivars likely originated from the donors with T3-type MKK3 in East Asia. Besides the cultivars with T2 and T3 MKK3, we also found highly nondormant cultivars with T1-type MKK3, and all these cultivars were collected from Ethiopia and the Fertile Crescent. Therefore, the cultivars in Ethiopia likely evolved from the third ancestral donors harboring the T1-type MKK3 (**Figure 7**). Genome-wide comparisons revealed three distinct MKK3 groups that are broadly present in wild barley accessions with the sequence divergence of three MKK3 gene groups predating barley domestication. We also estimated the divergence times of MKK3 genes belonging to the same groups between barley and wild progenitor and found that the divergence times far exceed the estimated timeframe of barley domestication (Data not shown). Thus, our findings support a model in which the wild progenitor diversified into three lineages, from which barley with different MKK3 haplotype groups were independently domesticated.

**Figure 7.**
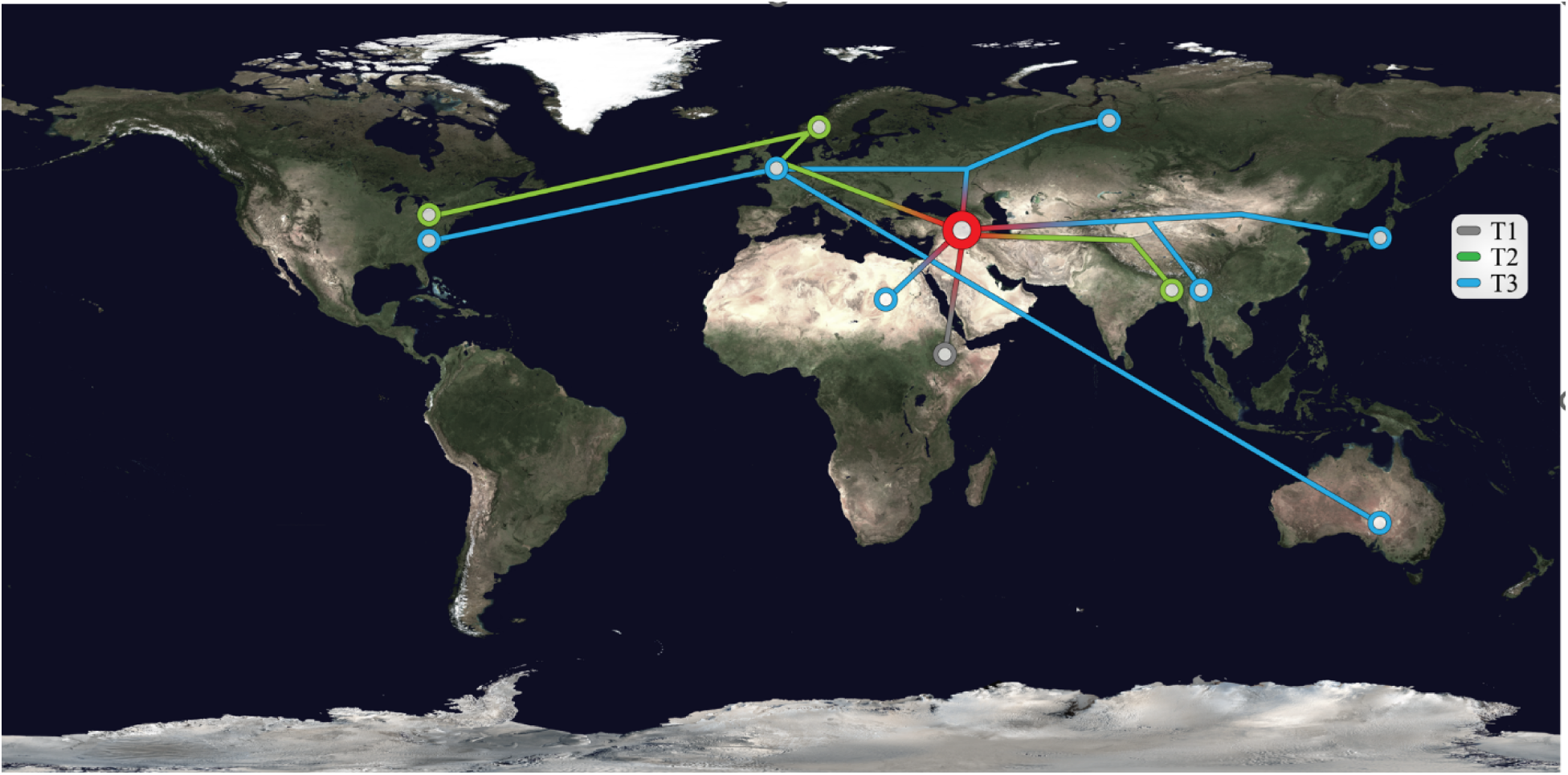
The proposed paths of highly germinated barley worldwide. Barley with three MKK3 groups was domesticated in the Fertile Crescent, the highly germinated barley varieties with T3-type MKK3 were widely spread across the world, the cultivars with T2-type MKK3 were distributed to China, Nepal, UK, Northern Europe and North America. The cultivars with T1-type MKK3 were only transferred to Ethiopia.

### Transposons drove MKK3 evolution

Growing evidence indicates that transposons play an integral role in genome evolution, providing the raw material for the birth of new genes (Volff 2006), reshaping gene expression, and driving phenotypic diversity including traits like grain size and seed shattering (Stude et al. 2011; Mao et al. 2022; Liu et al. 2022). Although over 80% of the barley genome consists of transposons and other repeats (Jayakodi et al. 2024), the role transposons play in shaping gene evolution is still ambiguous. Our findings provide compelling evidence that transposons within the barley genome have been a driving force behind MKK3 plasticity and enabled the phenotypic variation supporting adaptation to diverse environments fulfilling human needs. First, insertions/excision of MITEs introduced sequence variation within the MKK3 locus after the divergence of barley and bulbous barley. Second, similar transposon distributions were observed in the flanking regions of all duplicated MKK3 genes on chromosome 5H. While the mechanism underlying MKK3 CNV remains uncertain, transposons likely contributed to tandem amplification of MKK3 by providing ‘portable’ homologous regions for recombination or annealing (Reams and Roth 2015). Third, we also detected fragmentary MKK3 duplication resulting from gene capture and mobilization of a CACTA transposon. Several DNA transposon superfamilies including Mutator, Helitron and CACTA have been reported to capture gene fragments and drive gene shuffling, contributing to the emergence of novel gene functions (Jiang et al. 2004; Du et al. 2009; Catoni et al. 2019). However, the timing of these gene acquisition events has remained poorly understood. Our genomic comparison and evolutionary analysis indicate that the capture of five genes spanned at least 2.6 MY, with CAT250 acquiring fragments of three genes more than 4.5 MYA and incorporating the MKK3 fragment between 1.9-2.5 MYA, followed by a transposition. Beyond their critical role in MKK3 gene evolution, transposons also serve as valuable genomic markers for reconstructing the evolutionary history of MKK3.

In conclusion, this study reports the first comprehensive evidence describing the dynamic evolutionary history of MKK3 predating barley domestication and further underscores the critical role of transposons in shaping MKK3 diversity. Our findings reveal three independent origins of highly nondormant barley cultivars worldwide, including a unique lineage predominantly found in collections from Ethiopia. These results not only advance our understanding of the genomic basis of barley domestication but also offer a valuable foundation for breeding PHS tolerant cultivars optimized for diverse environments and production systems.

## MATERIALS AND METHODS

### Plant materials

43 cultivated and wild barley genotypes were collected from USDA seedbank or breeding programs, and the information on seed collections is shown in **Table S6**. The barley seeds were planted in the growth chamber at either the USDA - ARS Cereal Crops Research Unit (CCRU) in Madison, Wisconsin or the Small Grains and Potato Germplasm Research Unit (SGPGRU) in Aberdeen, Idaho. The young barley leaves were collected and used to extract genomic DNA using the cetyltrimethylammonium bromide (CTAB) method.

### MKK3 genes and sequence analysis

The 96 MKK3 gene sequences published by Nakamura et al. (2016) were downloaded from GenBank, and the accession numbers of all these genes started with “LC”. To identify the MKK3 genes in sequenced genomes of cultivated and wild barley, *Hordeum bulbosum* and wheat, the 8187-bp reference MKK3 gene was used to search against the NCBI genome datasets (https://www.ncbi.nlm.nih.gov/datasets/genome) and the best hits (with lowest E-values, highest sequence identities, and longest coverages) were extracted. All MKK3 genes previously published and identified in this study were used to compare with the reference MKK3 gene for determining the presence/absence of MITEs based on the sequence alignments and the structural features of MITEs including TIRs and TSDs.

### ddPCR and PCR analysis

dd PCR analysis was performed by following Collier et al. (2017). Briefly, four ug of barley genomic DNA for each sample was digested with the HindIII restriction enzyme (New England Biolabs, Ipswich, MA) at 37 °C for 1 h. The Master mixture was prepared with the ddPCR Supermix for Probes (No dUTP) (Bio-Rad Laboratories, Hercules, CA) by following the manufacturer’s protocol, the mixture (23 μl) contains 5 uM FAM probe (reference gene, Actin 1 (Gines et al. 2018) Probe: 5′-[6-FAM]TGTTTGAGACTTTCAATGTTCCTGCC[BHQ1a-Q]-3′), 18uM reference gene Primer F (Actin F1: 5′ CCCAAAAGCCAACAGAGAGA -3′), 18 uM reference gene Primer R (Actin R1: 5′-GCCTGAATAGCGACGTACAT-3′), 5uM HEX probe (probe includes 2,506^th^ nucleotide of reference MKK3 gene or the MKK3Q165 site, which is related to exchange of a Glutamic acid (E) to Glutamine (Q) at position 165 (Vetch et al. 2020) MKK3 SNP P6: 5′-[HEX] GCATTGCCCTT[G/C]AATACATGGAT [BHQ1a-Q]-3′), 18uM Primer F ( MKK3 SNP F2: 5′-TGAATTCCAGGGTGCATTTT -3′) and 18 uM Primer R (MKK3 SNP R6: 5′-ATATGTGCAAGAACCGGCTC-3′). The reaction mixture (25 μl) containing 250 ng digested template DNA and 22.5 μl Master mixture was added to a new 96-well PCR plate. PCR was conducted with a C1000 Touch™ Thermal Cycler (Bio-Rad Laboratories, Hercules, CA) under the following conditions: enzyme activation at 95 °C for 10 min, amplification of 60 cycles (94 °C for 30 s denaturation, 52 °C for 1 min annealing/extension), and enzyme deactivation at 98 °C for 10 min. Two independent reactions were run for each sample, and the ddPCR results were analyzed using the Bio-Rad QuantaSoft™ Analysis Pro software. Each of the two reactions were separately analyzed and their average represents the copy numbers of the reference gene and MKK3.

Two pairs of PCR primers were used to detect the presence/absence of the two MITEs in MKK3 gene including TE13F (5′-TGGAAATGGAGCAAGCAGTG -3′) and TE13R (5′-TTGTCCAGAATCAGGCATGT-3′) targeted the flanking regions of the 769-bp InDel including Hvu_MITE1a; TE22F (5′-GTCACCTGAGAGAATTCGTAA-3′) and TE22R (5′-AAGGAACAAAACTCTGGTGTA-3′) targeted the flanking regions of Hvu_MITE2. The designed primers were synthesized by the Eurofins Genomics LLC (Louisville, KY, United States) and applied for PCR analysis. PCR amplifications were conducted in 20 μl reactions consisting of ∼100 ng of genomic DNA, 0.2 mM primer, deionized water, and 10 μl Phusion High-Fidelity PCR Master Mix (New England Biolabs, Ipswich, MA) containing 0.4 units/μl of Phusion DNA Polymerase, Phusion HF Buffer, 200 μM dATP, 200 μM dGTP, 200 μM dCTP, 200 μM dATP, and 1.5 mM MgCl2. The PCR temperature cycling conditions were 1 cycle of 98°C for 30s; 35 cycles of 98°C for 10 s, 62°C for 15 s, 72°C for 30 s; and 1 cycle of 72°C for 5 min. Amplification products were run on 0.8% agarose gels and stained with ethidium bromide.

### Phylogenetic Tree Constructions

The complete sequences of 96 published MKK3 genes (Nakamura et al. 2016) and 86 MKK3 genes extracted from the sequenced barley genomes in this study were merged and used to conduct all against all comparisons. If multiple MKK3 genes showed 100% sequence identity between each other, only one sequence was included for phylogenetic analysis. After exclusion of the identical MKK3 genes, 179 MKK3 gene sequences were kept and aligned by MAFFT v7.490 (Katoh and Standley 2013) using Geneious Prime v.2025.2 (Kearse et al. 2012). The multiple sequence alignments were then analyzed with IQ-TREE 2 (Minh et al. 2020) using ModelFinder (Kalyaanamoorthy et al. 2017) to select the best-fitting ML model for phylogenetic analyses. Using the GTRGAMMA model selected by ModelFinder, sequence alignments were separately applied to build phylogenetic trees with RAxML software (Stamatakis 2014) using ML methods. The phylogenetic trees were uploaded to the Interactive Tree of Life website (Letunic and Bork 2021) and visualized.

### Estimation of MKK3 divergence times

All MKK3 genes were divided into three groups (T1-T3) based on the numbers of MITEs and used to conduct all against all sequence alignments. A custom Perl script developed (Yupeng Li, unpublished) was used to estimate the divergence among different groups using pairwise Kimura 2-parameter distance estimates calculated between all MKK3 sequence pairs. The divergence times were calculated using the formula: T = K/2r where K was kimura 2-parameter distance estimate and r was an average substitution rate of 6.5 x 10^9^ substitutions per synonymous site per year which is the molecular clock rate of the *Adh* gene in the grass family (Gaut et al. 1996). All statistical summaries and visualizations were performed in R using the tidyverse framework (Wickham et al. 2019). Divergence distributions were visualized using boxplots generated from ggplot2, with separate categories for each comparison between groups.

### Analysis of MKK3 expression

RNA-seq data of 20 barley accessions were published by the barley pan-transcriptome project (Guo et al. 2025), and all available biological replicates were included for this analysis. Transcript abundance was quantified using Salmon v.1.10.0 (Patro et al. 2017) with the default parameters. Reads were mapped to the PanBaRT20 barley pan-transcriptome reference, and transcript level expression estimates were reported as transcripts per million (TPM). Salmon output files (*quant.sf*) were generated for each sample and used for downstream analyses. Expression of the MKK3 genes was extracted based on the PanBaRT20 transcript annotation corresponding to MKK3 (PanBaRT20_chr5HG56507). TPM values for this transcript were extracted from each *quant.sf* file. Each barley accession was labeled based on MKK3 copy number (Jørgensen et al. 2025) and the number of MITEs. All statistical analyses were performed in R (R core Team 2025). Pairwise comparisons of MKK3 transcript abundance among different groups were conducted using Welch’s two-sample t-tests, which are robust to unequal variances and sample sizes. Nonparametric Wilcoxon rank-sum tests were also performed as a complementary approach. For visualization, transcript abundance values were averaged across biological replicates within each accession to generate a single mean TPM value per genotype. The relationship between MITE number and MKK3 transcript abundance was visualized using scatterplots and a linear regression line.

## Author Contributions

LT built the phylogenetic trees, analyzed the sequence data and gene expression and drafted the methods; AC extracted DNA and performed ddPCR and PCR analysis; JW initiated the project, conducted Sanger sequencing and drafted the manuscript; DG conceived the experiments, collected sequence data and drafted the manuscript. All authors were involved in manuscript revision.

## Acknowledgments

This research was supported by USDA-ARS CRIS Projects No. 5090-30600-001-000-D and 2050–21000-038-000D. We thank Dr. Ning Jiang for the valuable comments. Authors would like to express our gratitude to Leslie Zalapa for assisting with DNA isolations and Dr. Josh MacCready for his assistance with figures. We also thank Dr. Yupeng Li for sharing the script.

## Data Availability Statement

All relevant data are included in the manuscript and the supporting materials.

## References

Badr, A., Müller, K., Schäfer-Pregl, R., El Rabey, H., Effgen, S., Ibrahim, H. H., et al. (2000). On the origin and domestication history of barley (Hordeum vulgare). Mol. Biol. Evol. 17, 499–510.

Barrón, M.G., Fiston-Lavier, A.S., Petrov, D.A., & González, J. (2014) Population genomics of transposable elements in Drosophila. Annu Rev Genet. 48,561–81.

Beattie, A.D., Edney, M.J., Scoles, G.J., and Rossnagel, B.G. (2010) Association mapping of malting quality data from western Canadian two-row barley cooperative trials. Crop Sci 50(5), 1649–1663.

Bewley, J.D., Bradford, K.J., Hilhorst, H.W.M., Nonogaki, H. (2013). Seeds: Physiology of development, germination and dormancy. (3rd ed) Springer.

Brassac, J. & Blattner, F. R. (2015) Species-level phylogeny and polyploid relationships in *Hordeum* (Poaceae) inferred by next-generation sequencing and in silico cloning of multiple nuclear loci. Syst. Biol. 64, 792–808.

Bureau, T.E. & Wessler, S.R. (1992) Tourist – a large family of small inverted repeat elements frequently associated with maize genes. Plant Cell, 4, 1283–1294.

Catoni, M., Jonesman, T., Cerruti, E., & Paszkowski, J. (2019) Mobilization of Pack-CACTA transposons in Arabidopsis suggests the mechanism of gene shuffling. Nucleic Acids Res. 47(3), 1311–1320.

Clare, S.J., Alhashel, A.F., Li, M., Effertz, K.M., Poudel, R.S., Zhang, J. et al. (2024) High resolution mapping of a novel non-transgressive hybrid susceptibility locus in barley exploited by Pyrenophora teres f. maculata. BMC Plant Biology, 24, 622.

Collier, R., Dasgupta, K., Xing, Y.P., Hernandez, B.T., Shao, M., Rohozinski, D. et al. (2017) Accurate measurement of transgene copy number in crop plants using droplet digital PCR. Plant Journal, 90(5), 1014–1025.

Doebley, J.F., Gaut, B.S. & Smith, B.D. (2006) The molecular genetics of crop domestication. Cell, 127, 1309–1321.

Du, C., Fefelovam N., Caronnam, J., He, L., & Dooner, H.K. (2009) The polychromatic Helitron landscape of the maize genome. Proc Natl Acad Sci U S A. 106(47), 19916–21.

Fox, G.P., & Bettenhausen, H.M. (2023) Variation in quality of grains used in malting and brewing. Front. Plant Sci. 14,1172028.

Gaut, B., Morton, B., McCaig, B. & Clegg, M. (1996) Substitution rate comparisons between grasses and palms: synonymous rate differences at the nuclear gene Adh parallel rate differences at the plastid gene rbcL. Proceedings of the National Academy of Sciences, 93, 10274–10279.

Gines, M., Baldwin, T., Rashid, A., Bregitzer, P., Maughan, P.J., Jellen, E.N. et al. (2018) Selection of expression reference genes with demonstrated stability in barley among a diverse set of tissues and cultivars. Crop Sci, 58, 332–341.

Gubler, F., Millar, A.A., & Jacobson J.V. (2005) Dormancy release, ABA and pre-harvest sprouting Curr Opin Plant Biol. 8(2), 183–187.

Guo, W., Schreiber, M., Marosi, V., Bagnaresi, P., Chalmers, K.J., Chapman, B. et al. (2025) A barley pan-transcriptome reveals layers of genotype-dependent transcriptional complexity. Nature Genetics, 57, 441–450.

Han, F., Ullrich, S.E., Clancy J.A., Jitkov, V., Kilian A., & Romagosa, I. (1996) Verification of seed dormancy loci via linked molecular markers. Theor and Appl Genet, 92(1), 87–91.

Han, Y. & Wessler, S.R. (2010) MITE-Hunter: a program for discovering miniature inverted-repeat transposable elements from genomic sequences. Nucleic Acids Res. 38, e199.

Hu, H., Wang, P., Angessa, T.T., Zhang, X.-Q., Chalmers, K.J., Zhou, G. et al. (2023) Genomic signatures of barley breeding for environmental adaptation to the new continents. Plant Biotechnology Journal, 21, 1435–1448.

Jayakodi, M., Lu, Q., Pidon, H., Rabanus-Wallace, M.T., Bayer, M., Lux, T. et al. (2024) Structural variation in the pangenome of wild and domesticated barley. Nature, 636, 654–666.

Jiang. N., Bao, Z., Zhang, X., Eddy, S.R., & Wessler, S.R. (2004) Pack-MULE transposable elements mediate gene evolution in plants. Nature. 431(7008):569–73.

Jiang, C., Lei, M., Guo, Y., Gao, G., Shi, L., Jin, Y. et al. (2022) A reference-guided TILLING by amplicon-sequencing platform supports forward and reverse genetics in barley. Plant Communications, 3, 100317.

Jørgensen, M.E., Vequaud, D., Wang, Y., Andersen, C.B., Bayer, M., Box, A., et al. (2026) Postdomestication selection of *MKK3* shaped seed dormancy and end-use traits in barley. Science. 391, 90–95.

Kalyaanamoorthy, S., Minh, B.Q. Wong, T.K.F., von Haeseler, A., Jermiin, L.S. (2017) ModelFinder: fast model selection for accurate phylogenetic estimates. Nat Methods. 14:587–589.

Katoh, K., & Standley, D.M. (2013) MAFFT multiple sequence alignment software version 7: improvements in performance and usability. Mol Biol Evol. 30:772–780.

Kearse, M., Moir, R., Wilson, A., Stones-Havas, S., Cheung, M., Sturrock, S., et al. (2012) Geneious Basic: an integrated and extendable desktop software platform for the organization and analysis of sequence data. Bioinformatics. 28:1647–1649.

Kjær K.H., Ruter, A.H., Mendendez-Serra, M., Vogel, N.A., Ramsøe, A.D., Farnsworth, W.R., et al. (2025) Environmental DNA reveals reykjavik’s human and ecological history. bioRxiv. 10.08.681091.

Le Rouzic, A., Payen, T., & Hua-Van, A. (2013) Reconstructing the evolutionary history of transposable elements. Genome Biol Evol. 5(1), 77–86.

Letunic, I., & Bork, P. (2024) Interactive Tree of Life (iTOL) v6: recent updates to the phylogenetic tree display and annotation tool. Nucleic Acids Res. 52(W1):W78–W82.

Lin, R., Horsely, R.D., Lapitan, N.L.V, Ma, Z., & Schwarz, P.B. (2009). WTL mapping of dormancy in barley using the Harrington/Morex and Chevron/Stander populations. Crop Sci, 49(3), 841–849.

Liu, H., Fang, X., Zhou, L., Li, Y., Zhu, C., Liu, J. et al. (2022) Transposon insertion drove the loss of natural seed shattering during Foxtail Millet domestication. Molecular Biology and Evolution. 39, msac078.

Liu, M., Li, Y., Ma, Y., Zhao, Q., Stiller, J., Feng, Q., et al. (2020) The draft genome of a wild barley genotype reveals its enrichment in genes related to biotic and abiotic stresses compared to cultivated barley. Plant Biotechnology Journal, 18, 443–456.

Lynch, M., & Conery, J.S. (2000) The evolutionary fate and consequences of duplicate genes. Science. 290(5494):1151–5.

Mao, D., Tao, S., Li, X., Gao, D., Tang, M., Liu, C. et al. (2022) The harbinger transposon-derived gene PANDA epigenetically coordinates panicle number and grain size in rice. Plant Biotechnology Journal, 20(6), 1154–1166.

Matilla, A. J. (2024). Current insights into weak seed dormancy and pre-harvest sprouting in crop species. Plants 13: 2559.

Minh, B.Q., Schmidt. H.A., Chernomor. O., Schrempf. D., Woodhams. M.D., von Haeseler, A., et al. (2020) IQ-TREE 2: new models and efficient methods for phylogenetic inference in the genomic era. Mol Biol Evol. 37:1530–1534.

Nagel, M., Alqudah, A.M., Bailly, M., et al. (2019) Novel loci and a role for nitric oxide for seed dormancy and preharvest sprouting in barley. Plant Cell Environ. 42(4), 1318–1327.

Nakamura, S., Pourkheirandish, M., Morishige, H. et al. (2016) Mitogen-activated protein kinase kinase 3 regulates seed dormancy in barley. Curr. Biol. 26, 775–781.

Pan, R., Hu, H., Xiao, Y., Xu, L., Xu, Y., Ouyang, K. et al. (2023) High-quality wild barley genome assemblies and annotation with Nanopore long reads and Hi-C sequencing data. Scientific Data, 10, 535.

Patro, R., Duggal, G., Love, M.I., Irizarry, R.A., Kingsford, C. (2017) Salmon provides fast and bias-aware quantification of transcript expression. Nature Methods. 14:417–419.

Qian, W., Liao, B.Y., Chang, A.Y., & Zhang, J. (2010) Maintenance of duplicate genes and their functional redundancy by reduced expression. Trends Genet. 26(10):425–30.

R Core Team. (2025). R: A Language and Environment for Statistical Computing. R Foundation for Statistical Computing, Vienna, Austria. https://www.R-project.org

Reams, A.B., & Roth, J.R. (2015) Mechanisms of gene duplication and amplification. Cold Spring Harb Perspect Biol. 7(2), a016592.

Rodriguez, M.V., Barrero, J.M., Corbineau, F., Gubler, F., & Benech-Arnold, R.L. (2015) Dormancy in cereals (not too much, not so little): About the mechanisms behind this trait. Seed Science Research, 25(2), 99–119.

Rooney, T.E., Sweeney, D.W., Kunze, K.H., Sorrells, M.E., and Walling, J.G. (2023) Malting quality and preharvest sprouting are genetically correlated in spring malting barley. Theor Appl Genet. 136(3), 59.

Sato, K., Yamane, M., Yamaji, N. et al. (2016) Alanine aminotransferase controls seed dormancy in barley. Nat Commun 7, 11625 (2016).

Scott, L., LaFoe, D. and Weil, C.F. (1996) Adjacent sequences influence DNA repair accompanying transposon excision in maize. Genetics, 29, 237–246.

Stamatakis, A. (2014) RAxML version 8: a tool for phylogenetic analysis and post-analysis of large phylogenies. Bioinformatics. 30(9):1312–1313.

Studer, A., Zhao, Q., Ross-Ibarra, J. & Doebley, J. (2011) Identification of a functional transposon insertion in the maize domestication gene tb1. Nat. Genet. 43, 1160–1163.

Sweeney, D.W., Rooney, T.E., Walling, J.G. & Sorrells, M.E. (2021). Interaction of the barley SD1 and SD2 seed dormancy loci influence preharvest sprouting, seed dormancy, and malting quality. Crop Sci, 62, 120–138.

Ullrich, S.E., Lee, H., Clancy, J.A. et al. (2009) Genetic relationships between preharvest sprouting and dormancy in barley. Euphytica, 168, 331–345.

Vetch, J.M., Walling, J.G., Sherman, J., Martin, J.M. and Giroux, M.J. (2020) Mutations in the *HvMKK3* and *HvAlaAT1* genes affect barley preharvest sprouting and after-ripened seed dormancy. Crop Sci. 60, 1897–1906.

Volff, J.N. (2006) Turning junk into gold: Domestication of transposable elements and the creation of new genes in eukaryotes. BioEssays. 28, 913–922.

Wicker, T., Sabot, F., Hua-Van, A., Bennetzen, J.L., Capy, P., Chalhoub, B. et al. (2007) A unified classification system for eukaryotic transposable elements. Nature Reviews Genetics, 8, 973–982.

Wickham, H., Averick, M., Bryan, J., Chang, W., McGowan, L., François, R., et al. (2019). Welcome to the tidyverse. Journal of Open Source Software. 4(43): 1686.

Wong, L.H., & Choo, K.H. (2014) Evolutionary dynamics of transposable elements at the centromere. Trends Genet. 20(12), 611–6.

Woonton, B.W., Jacobsen, J.V., Sherkat, F., & Stuart I.M. (2005). Changes in germination and malting quality during storage of barley. J. Inst Brewing, 111(1), 33–41.

Xu, W., Tucker, J.R., Bekele, W.A., You, F.M., Fu, Y.-B., Khanal, R. et al. (2021) Genome assembly of the Canadian two-row malting barley cultivar AAC Synergy. G3: Genes, Genomes, Genetics, 11, jkab031.

Zeng, X., Xu, T., Ling, Z., Wang, Y., Li, X., Xu, S. et al. (2020) An improved high-quality genome assembly and annotation of Tibetan hulless barley. Scientific Data, 7, 139.

Zhang, M., Pan, J., Kong, X., Zhou, Y., Liu, Y., Sun, L., & Li, D. (2012). ZmMKK3, a novel maize group B mitogen-activated protein kinase kinase gene mediates osmotic stress and ABA signal responses. J. Plant Physiol. 169, 1501–1510

