## Supplementary Figs for "Transposons Triggered Dynamic Evolution of MKK3 Gene, a Key Regulator for Seed Dormancy in Barley"

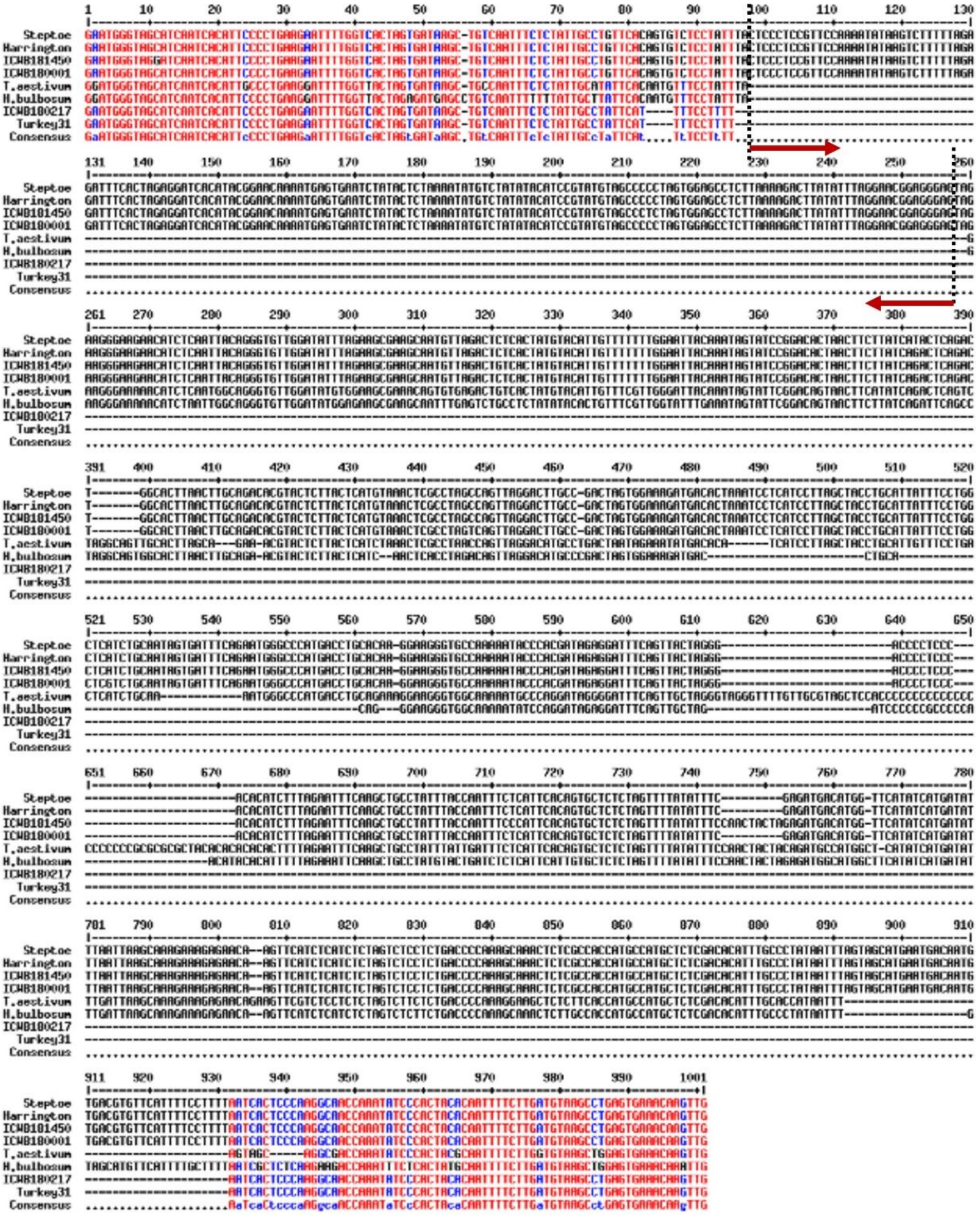
Steptoe
Harrington
ICWB181450
ICWB180001
T.aestivum
H.bulbosum
ICWB180217
Turkey31
Consensus

A

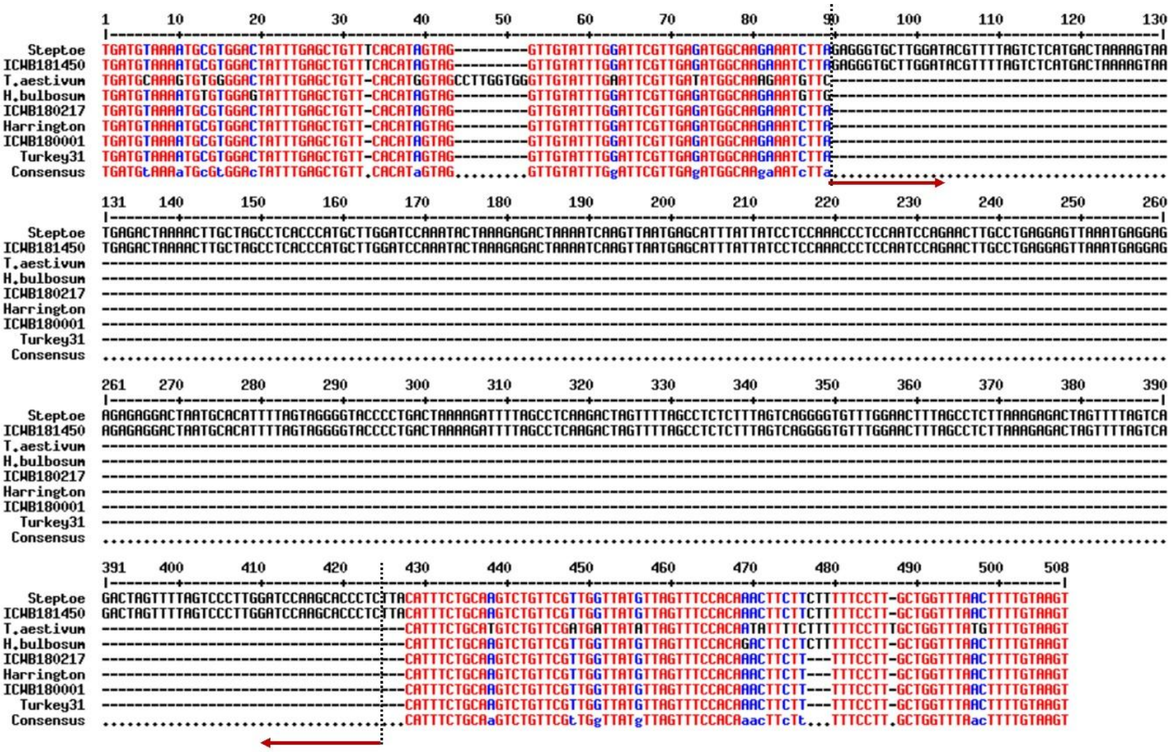

B

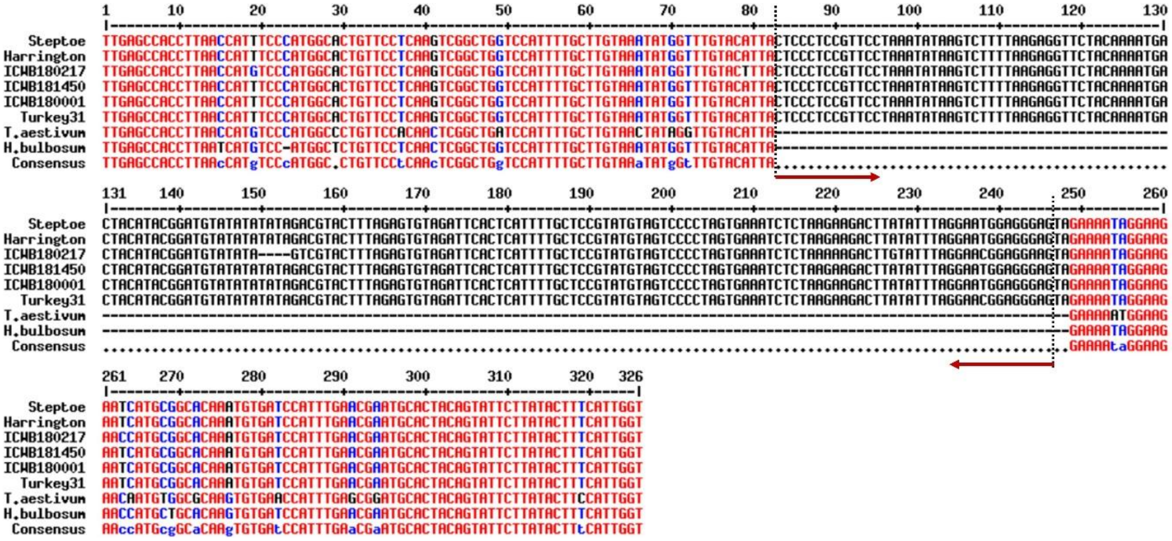

C

**Figure S1. Alignment of the 769-bp InDel (A), 338-bp InDel (B) and Hvu_MITE1b (C) and their 200-bp flanking sequences (100-bp for each side) in MKK3 gene from six cultivated and wild barley, bulbous barley and wheat**. The vertical broken lines indicated the boundaries of MITEs, and the the horizontal lines with arrow indicated the 13-bp TIRs (CTCCCTCCGTTCC… GGAACGGAGGGAG) for Hvu_MITE1 and 14-bp TIRs (GAGGGTGCTTGGAT… ATCCAAGCACCCTC) for Hvu_MITE2.

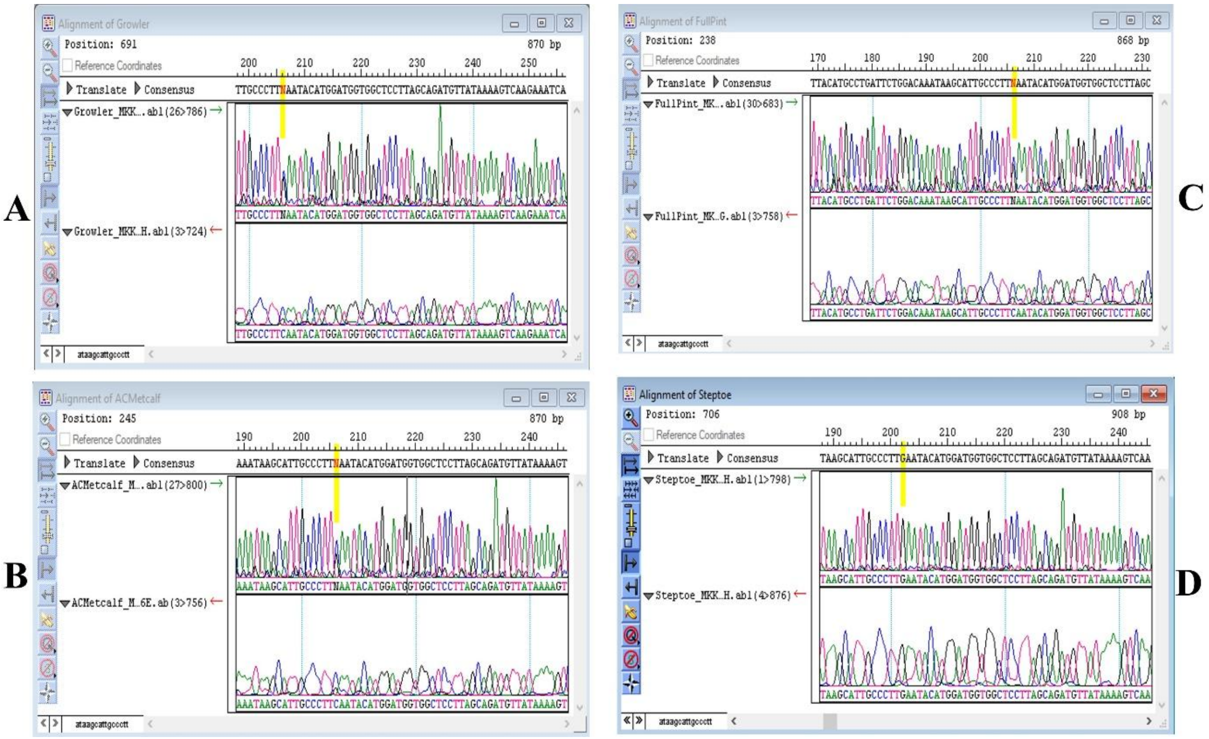


**Figure S2. Sanger sequencing of MKK3 gene around E165Q in Growler (A), AC Metcalfe (B), Full Pint (C) and Steptoe (D).** Double peaks were observed at E165Q in Growler, AC Metcalfe and Full Pint.

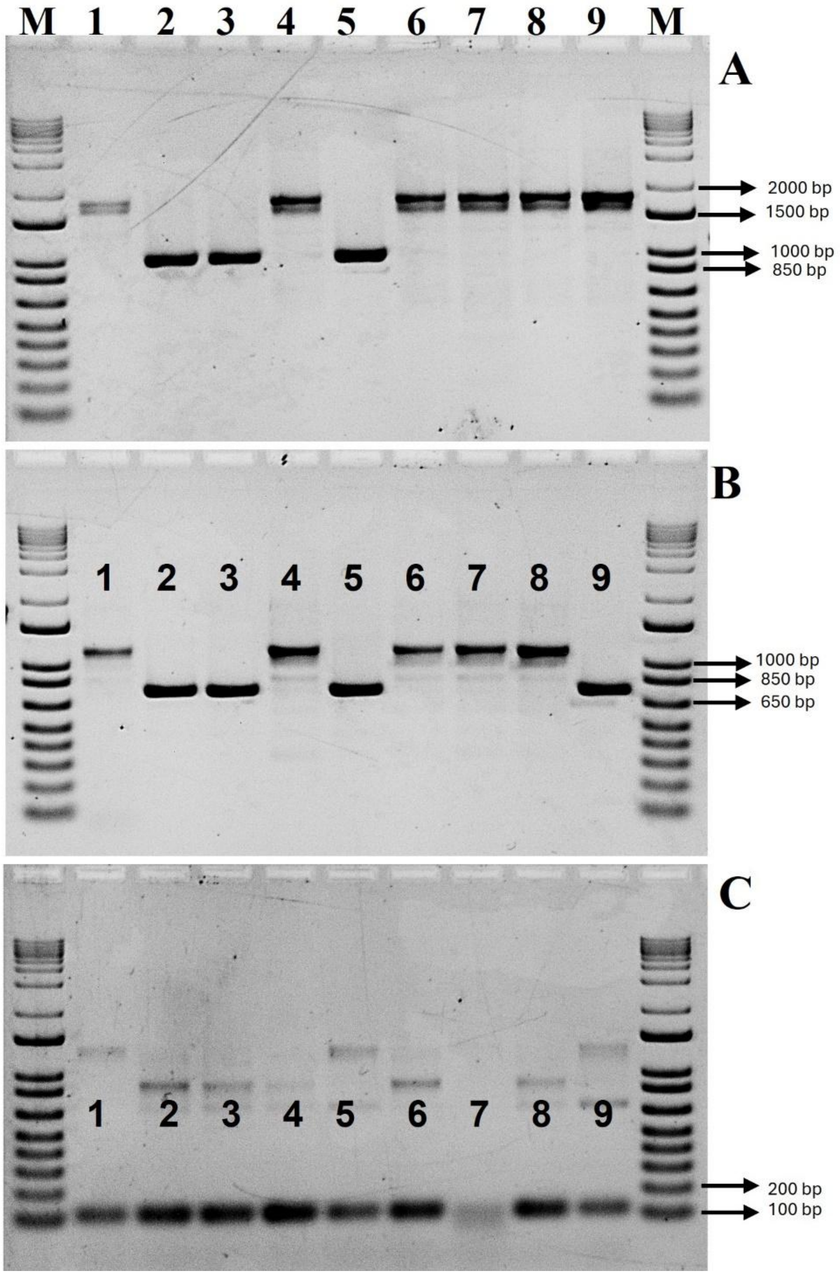


**Figure S3. PCR analysis with the primers for amplifying the flanking regions of 769-bp InDel containing Hvu_MITE1a (A), Hvu_MITE2 (B) and Actin gene (C).** The expected size was 941 bp (without Hvu_MITE1a) and 1710-bp (with Hvu_MITE1a) for A, and 741-bp (without Hvu_MITE2) and 1079-bp (with Hvu_MITE2) for B. 1: PI681799, 2: PI681815, 3: PI681843, 4: PI 681890, 5: PI 681905, 6: PI 681942, 7: Golden Promise, 8: Morex. M: 1-Kb Plus DNA ladder.

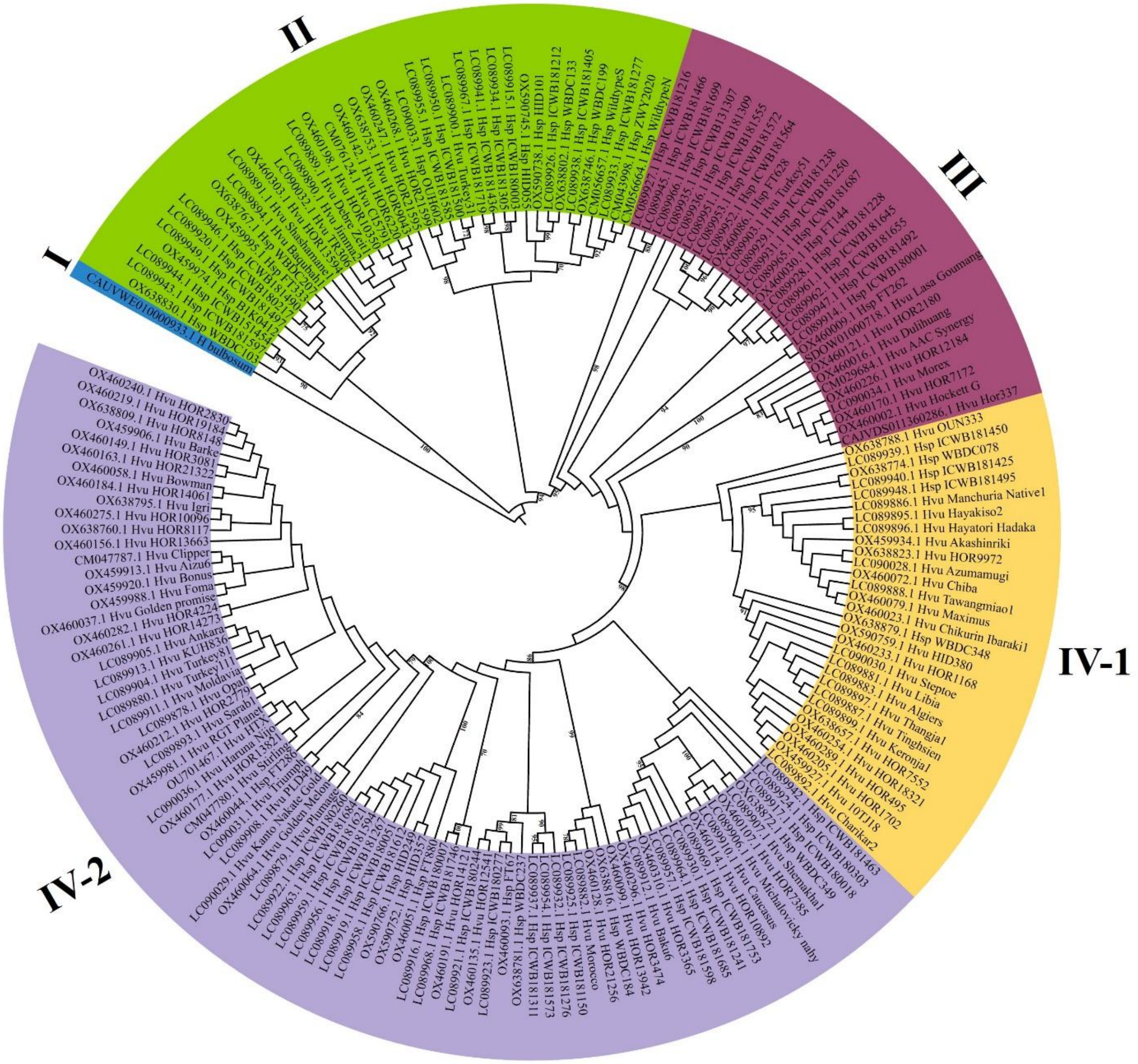


**Figure S4. Phylogenetic tree of 179 MKK3 gene sequences removing the two InDels regions in cultivated and wild barley built with maximum likelihood (ML) method**. The bootstrap values of ＞70% are labeled. The three letters between the GenBank accession number and the cultivar (wild accession) represent cultivated barley (Hvu) and wild progenitor (Hsp), respectively.
